# Genetic background and mistranslation frequency determine the impact of mistranslating tRNA^Ser^_UGG_

**DOI:** 10.1101/2022.03.22.485345

**Authors:** Matthew D. Berg, Yanrui Zhu, Raphaël Loll-Krippleber, Bryan-Joseph San Luis, Julie Genereaux, Charles Boone, Judit Villen, Grant W. Brown, Christopher J. Brandl

## Abstract

Transfer RNA variants increase the frequency of mistranslation, the mis-incorporation of an amino acid not specified by the “standard” genetic code, to frequencies approaching 10% in yeast and bacteria. Cells cope with these variants by having multiple copies of each tRNA isodecoder and through pathways that deal with proteotoxic stress. In this study, we define the genetic interactions of the gene encoding tRNA^Ser^_UGG,G26A_, which mistranslates serine at proline codons. Using a collection of yeast temperature sensitive alleles, we identify negative synthetic genetic interactions between the mistranslating tRNA and 109 alleles representing 91 genes, with nearly half of the genes having roles in RNA processing or protein folding and turnover. By regulating tRNA expression, we then compare the strength of the negative genetic interaction for a subset of identified alleles under differing amounts of mistranslation. The frequency of mistranslation correlated with impact on cell growth for all strains analyzed; however, there were notable differences in the extent of the synthetic interaction at different frequencies of mistranslation depending on the genetic background. For many of the strains the extent of the negative interaction with tRNA^Ser^_UGG,G26A_ was proportional to the frequency of mistranslation or only observed at intermediate or high frequencies. For others the synthetic interaction was approximately equivalent at all frequencies of mistranslation. As humans contain similar mistranslating tRNAs these results are important when analyzing the impact of tRNA variants on disease, where both the individual’s genetic background and the expression of the mistranslating tRNA variant need to be considered.

## INTRODUCTION

Mistranslation occurs when an amino acid that differs from what is specified by the standard genetic code is incorporated into a growing polypeptide chain during translation. Mistranslation occurs in all cells but can be enhanced by environmental conditions or mutations in the translational machinery (Lee *et al*. 2006; Netzer *et al*. 2009; Ling and Söll 2010; Jones *et al*. 2011; Wiltrout *et al*. 2012; Reverendo *et al*. 2014; Lant *et al*. 2017; Schwartz and Pan 2017). Mutations in tRNAs that cause mistranslation were initially identified as intergenic suppressors that change the meaning of the genetic code (Stadler and Yanofsky 1959; Yanofsky and Crawford 1959; Crawford and Yanofsky 1959; Benzer and Champe 1962; Gorini and Beckwith 1966). tRNA^Ser^ variants are particularly prone to mistranslate because the anticodon is not a major identity element for aminoacylation by the cognate serine aminoacyl-tRNA synthetase (Giegé *et al*. 1998). Rather, specificity for aminoacylation comes from the long variable arm positioned 3’ of the anticodon stem (Asahara *et al*. 1994; Biou *et al*. 1994; Himeno *et al*. 1997). Therefore, anticodon mutations in tRNA^Ser^ encoding genes lead to mis-incorporation of serine at non-serine codons (Geslain *et al*. 2010; Berg *et al*. 2017, 2019b; Zimmerman *et al*. 2018). Interestingly, human genomes contain similar tRNA^Ser^ variants and other variant tRNAs with the potential to mistranslate (Berg *et al*. 2019a; Lant *et al*. 2019). In zebrafish and flies, mistranslating tRNA variants reduce viability and increase the frequency of deformities (Reverendo *et al*. 2014; Isaacson *et al*. 2022).

The toxic effects of mistranslating tRNAs are buffered through multiple copies of each tRNA isodecoder (for example, there are 275 tRNA-encoding genes in *Saccharomyces cerevisiae;*Chan and Lowe 2016) and through protein quality control mechanisms that deal with misfolded protein and protein aggregates (reviewed in Hoffman *et al*. 2017). When mistranslation reaches a threshold, protein quality control mechanisms no longer protect the cell and growth is impaired (Berg *et al*. 2019b). The extent of growth impairment is inversely related to the frequency of mistranslation in a linear fashion with yeast growth being arrested when mistranslation approaches approximately 12% (Berg *et al*. 2021a).

We previously demonstrated that the negative genetic interactions with mistranslating tRNAs depend on the amino acid substitution (Berg *et al*. 2021b). At similar frequencies of mistranslation, a tRNA variant substituting serine at arginine codons has more genetic interactions than one substituting alanine at proline codons. In this report we identify the genetic interactions of a mistranslating serine tRNA variant that incorporates serine at proline codons. Using a regulated tRNA expression system, we show that although there is a general correlation between the frequency of mistranslation and impact on growth, the impact of different mistranslation frequencies depends on a strain’s specific genetic background. As similar mistranslating tRNAs are found in the human population, these results suggest that genetic background contributes to the impact of tRNA variants on health and disease.

## MATERIALS AND METHODS

### Yeast strains and growth

BY4742 (*MATα his3Δ0 leu2Δ0 lys2Δ0 ura3Δ0*; Brachmann *et al*. 1998) and Y7092 (*MATα can1Δ*::*STE2pr-SpHIS5 lyp1Δ his3Δ1 leu2Δ0 ura3Δ0 met15Δ0*) strains are derivatives of S288c. Y7092 was a kind gift from Dr. Brenda Andrews (University of Toronto). Strains from the temperature sensitive collection are derived from the wild-type *MAT***a** haploid yeast strain BY4741 and described in Costanzo *et al*. (2016). The strains containing the gene expressing tRNA^Ser^_UGG, G26A_ (CY8613) were made by integrating modified *SUP17* and flanking sequence into Y7092 at the *HO* locus and selecting for the *natMX* marker, as previously described in Zhu *et al*. (2020), using the construct described below. The control strain (CY8611) was made by integrating only the *natNT2* marker at the *HO* locus. Transformants were selected on 100 μg/mL nourseothricin-dihydrogen sulfate (clonNAT) and integration was verified by PCR.

Yeast strains were grown at 30° in yeast peptone media containing 2% glucose (YPD) or synthetic media supplemented with nitrogen base and amino acids, unless otherwise indicated. Growth curves were generated by diluting saturated cultures to OD_600_ = 0.1 in synthetic complete media and incubating at 30°. OD_600_ was measured every 15 minutes for 24 hr using a BioTek Epoch 2 microplate spectrophotometer. Doubling time was calculated using the R package “growthcurver” (Sprouffske and Wagner 2016).

### DNA constructs

The construct to integrate the gene encoding tRNA^Ser^_UGG, G26A_ at the *HO* locus was created using a synthetic DNA containing 200 bp up and downstream of the *HO* translational start, previously described in Zhu *et al*. (2020). The construct was cloned into pGEM^®^-T Easy (Promega Corp.) as a *Not*I fragment to create pCB4386. The *natNT2* marker from pFA6-natNT2 was PCR amplified using primers UK9789/UK9790 (Table S1) and cloned into pCB4386 as an *Eco*RI fragment to generate the control SGA integrating vector (pCB4394). The gene encoding tRNA^Ser^_UGG,G26A_ was PCR amplified from pCB4023 (Berg *et al*. 2017) using primers UG5953/VB2609 and cloned as *Hin*dIII fragments into pCB4394 to create pCB4397.

*URA3*-containing centromeric plasmids expressing tRNA^Ser^ (pCB3076), tRNA^Ser^_UGG,G26A_ (pCB4023), tRNA^Ser^_UGG,G26A_ with 5’ *GAL1pr* (pCB4568), tRNA^Ser^_UGG,G26A_ with 3’ *GAL1pr* (pCB4566) are described in Berg *et al*. (2017, 2021a).

### Synthetic genetic array analysis and validation

The SGA assay was performed as described by Tong *et al*. (2001) with minor modifications. Strains CY8611 (*HO::natMX*) and CY8613 (*HO:: tRNA^Ser^_UGG, G26A_-natMX*) were crossed to a yeast temperature sensitive collection (Ben-Aroya *et al*. 2008; Li *et al*. 2011; Kofoed *et al*.2015; Costanzo *et al*. 2016) in quadruplicate 1536 colony array format using a BioMatrix (S&P Robotics Inc.) automated pinning system. In this format, each allele of the temperature sensitive collection is present in technical quadruplicate on the plate. Double mutants were selected on YPD plates containing 200 mg/L G418 and 100 mg/L clonNAT. Diploids were sporulated on enriched sporulation media and *MAT***a** haploid double-mutants selected using standard SGA media. The entire SGA procedure was carried out at room temperature, except for the colony scoring in order to minimize growth defect of the temperature sensitive strains. To identify genetic interactions, double mutants were pinned onto double mutant selection SGA medium and grown at 30° for 5 days. Images were taken every 24 hours to determine colony size computationally. SGATools (Wagih *et al*. 2013) was used to determine genetic interaction scores using a multiplicative model (ε = W_AB_ – W_A_ * W_B_; where ε is the interaction score, *W_AB_* is the fitness of the double mutant and *W_A_* and *W_B_* are the fitness values of each single mutant). Double mutant strains with an average interaction score less than −0.2 and Benjamini-Hochberg corrected *p*-value less than 0.05 were considered as potential negative genetic interactions.

Double mutants that were identified as negative genetic interactions from the screen were validated by re-creating the double mutant strain, starting from the single mutant haploid strains, using the SGA approach. Double mutant strains were grown in liquid media to saturation, cell densities were normalized, and cultures were spotted on SGA media. The temperature sensitive mutant crossed with the control strain CY8611 and the mistranslating tRNA strain crossed with a control *his3Δ* strain were also spotted to determine fitness of the single mutants. Intensity of each spot was measured with ImageJ (Schneider *et al*. 2012). Expected double mutant growth was calculated based on the growth of the single mutants and compared to the experimental growth of the double mutant. Double mutants that grew more slowly than expected were considered validated negative genetic interactions. Raw and validated data can be found in supplemental file S2.

Synthetic interactions with various frequencies of mistranslation were assessed by transforming the relevant temperature sensitive strains with *URA3*-containing centromeric plasmids expressing tRNA^Ser^ (pCB3076), tRNA^Ser^_UGG,G26A_ (pCB4023), tRNA^Ser^_UGG,G26A_ with 5’ *GAL1pr* (pCB4568) or tRNA^Ser^_UGG,G26A_ with 3’ *GAL1pr* (pCB4566). At least three individual transformants for each plasmid and strain were grown in synthetic complete medium lacking uracil and containing 2% galactose as the carbon source. Cells were grown to confluency, diluted 33-fold in 1x yeast nitrogen base and 5 μL was spotted onto solid media lacking uracil and containing 2% galactose. Cells were grown at 30° for 32-56 hours, depending on the strain, to achieve a level of growth (for the strain without mistranslation) consistent with the wild-type. Mean density of growth was determined with ImageJ (Schneider *et al*. 2012) and normalized growth for each mistranslating tRNA was calculated as a percent of the wild-type tRNA^Ser^ containing strain.

### SAFE analysis

Spatial analysis of functional enrichment (SAFE; Baryshnikova 2016) analysis was performed through TheCellMap (http://thecellmap.org; Usaj *et al*. 2017).

### Heat shock assay

Yeast strains containing the *HSE-GFP* reporter and a mistranslating tRNA variant were grown to stationary phase in medium lacking uracil and containing 0.6% casamino acids, diluted 1:100 in the same medium and grown for 18 hr at 30°. Cell densities were normalized to OD_600_ before measuring fluorescence with a BioTek Synergy H1 microplate reader at an excitation wavelength of 488 nm and emission wavelength of 528 nm. The mean relative fluorescence units were calculated from three technical replicates for each biological replicate.

### Mass spectrometry

Liquid chromatography tandem mass spectrometry to identify mistranslation was performed on five biological replicates of each strain. Starter cultures of each strain were grown overnight in YPD before being diluted to an OD_600_ of 0.1 in the same media and grown to an OD_600_ of ~ 1.0. Cells were lysed in a urea lysis buffer (8 M Urea, 50 mM Tris pH 8.2, 75 mM NaCl) and proteins were reduced with dithiothreitol and alkylated with iodoacetamide. Robotic purification and digestion of proteins into peptides was performed on the KingFisher^™^ Flex using LysC and the R2-P1 method described in Leutert *et al*. (2019).

Peptides were analyzed on a hybrid quadrupole orbitrap mass spectrometry (Orbitrap Exploris 480; Thermo Fisher Scientific) equipped with an Easy1200 nanoLC system (Thermo Fisher Scientific) as previously described in Berg *et al*. (2021b).

MS/MS spectra were searched against the *S. cerevisiae* protein sequence database (downloaded from the Saccharomyces Genome Database resource in 2014) using Comet (release 2015.01; Eng *et al*. 2013). The precursor mass tolerance was set to 50 ppm. Constant modification of cysteine carbamidomethylation (57.0215 Da) and variable modification of methionine oxidation (15.9949 Da) and proline to serine substitution (−10.0207 Da) were used for all searches. A maximum of two of each variable modification were allowed per peptide. Search results were filtered to a 1% false discovery rate at the peptide spectrum match level using Percolator (Käll *et al*. 2007). The mistranslation frequency was calculated using the unique mistranslated peptides for which the non-mistranslated sibling peptide was also observed. The frequency is defined as the counts of mistranslated peptides, where serine was inserted for proline, divided by the counts of all peptides containing proline and expressed as a percentage.

### Data availability

Strains and plasmids are available upon request. The authors affirm that all data necessary for confirming the conclusions of the article are present within the article, figures, and supplemental material. Supplemental File S1 contains all supplemental figures. Supplemental File S2 contains raw and validated SGA data. The mass spectrometry proteomics data have been deposited to the ProteomeXchange Consortium via the PRIDE (Perez-Riverol *et al*. 2019) partner repository with the dataset identifiers PDX025934 and PXD032063.

## RESULTS AND DISCUSSION

### Synthetic genetic interactions with tRNA^Ser^_UGG,G26A_

Yeast cells expressing tRNA^Ser^_UGG,G26A_, which contains a proline anticodon, mistranslate serine at proline codons (Berg *et al*. 2019b, 2021a). The G26A mutation is required in combination with the anticodon change to dampen otherwise lethal levels of mistranslation (Berg *et al*. 2017). To perform the SGA analysis, the gene encoding tRNA^Ser^_UGG,G26A_ (Figure 1A), including approximately 300 base pairs of 5’ and 3’ flanking sequence and a clonNAT resistance marker, was integrated at the *HO* locus. A control strain was created with only the clonNAT resistance marker integrated at the *HO* locus. Mass spectrometry-based analysis of the cellular proteome identified 4.9% proline to serine substitution in the strain expressing tRNA^Ser^_UGG,G26A_ compared to only 0.6% substitution in the control strain (Figure 1B). As shown in Figures 1C and 1D, tRNA^Ser^_UGG,G26A_ reduces cell growth (doubling time of 98 minutes versus 84 for the control strain) and results in a heat shock response (6.4-fold greater than the control strain).

**Figure 1.**
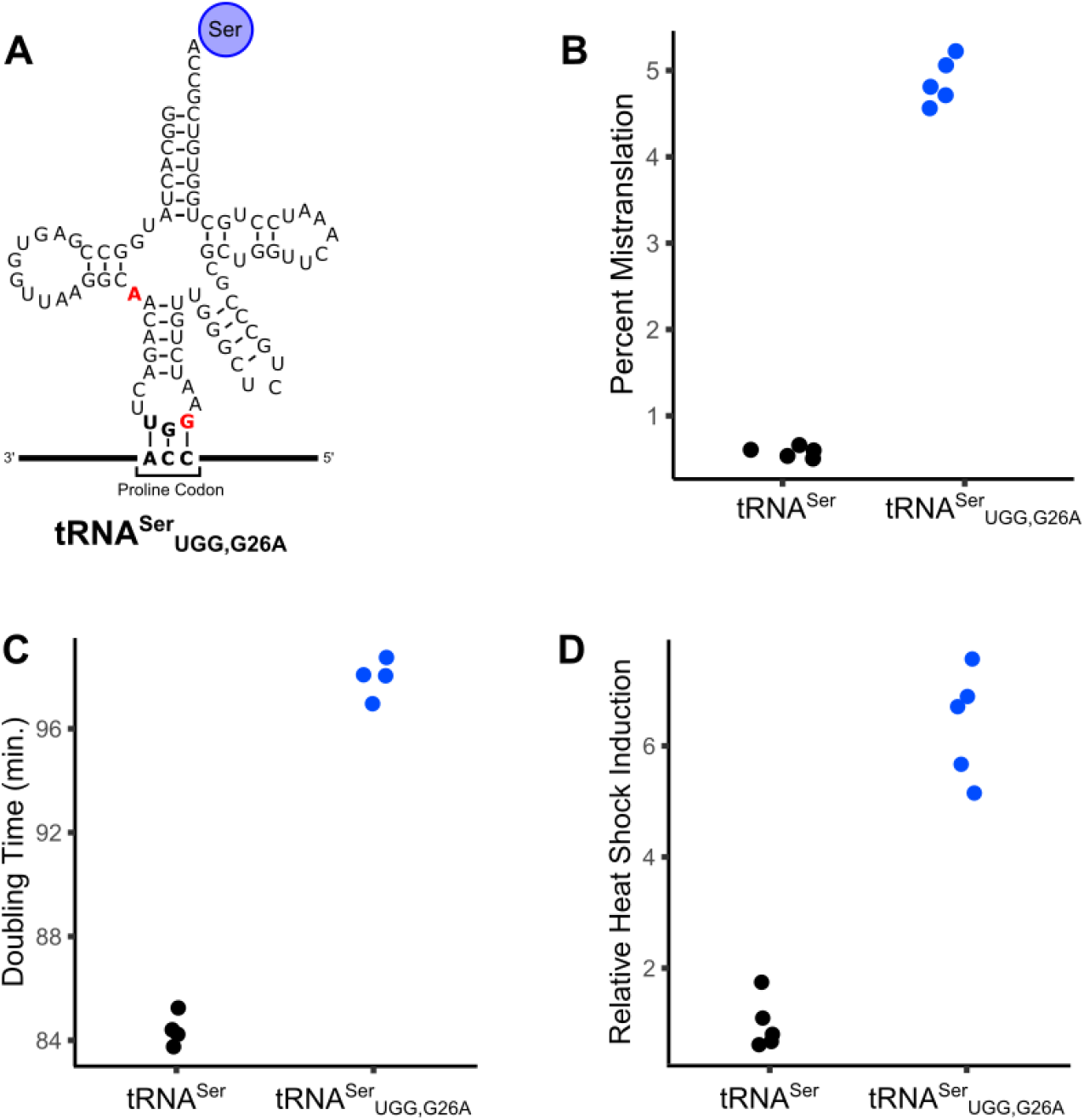
Phenotypic characterization of mistranslating tRNA^Ser^_UGG,G26A_. **A.** Cloverleaf representation of tRNA^Ser^_UGG,G26A_. The anticodon and G26 substitutions from the wild-type tRNA^Ser^ are shown in red. **B.** Mass spectrometry based analysis of the cellular proteome was performed on the control strain with no additional tRNA (CY8611) and the strain expressing mistranslating tRNA^Ser^_UGG,G26A_ (CY8613). Mistranslation frequency was calculated from the number of unique mistranslated peptides for which the non-mistranslated sibling peptide was also observed. Frequency is defined as the counts of peptides with serine substituted for proline divided by all peptides containing proline and expressed as a percentage. Each point represents one biological replicate (n = 5). Mistranslation frequency in the strain expressing tRNA^Ser^_UGG,G26A_ is statistically different compared to the control strain (Welch’s *t*-test; Bonferroni corrected *p*-value < 0.05). **C.** Doubling times for the strains described in B were determined from growth curves of the strains diluted to an OD_600_ ~ 0.1 in synthetic complete media containing clonNAT and grown for 24 hr. Doubling time was calculated with the R package “growthcurver” (Sprouffske and Wagner 2016). Each point represents one biological replicate (n = 4). Doubling time is statistically different between the strain expressing tRNA^Ser^_UGG,G26A_ and the control strain (Welch’s *t-*test; Bonferroni corrected *p-*value < 0.05). **D.** Strains described in B were transformed with a GFP reporter transcribed from a promoter containing heat shock response elements, grown to saturation in media lacking uracil, diluted 1:100 in the same media and grown for 18 hours at 30°. Cell densities were normalized and fluorescence measured. Each point represents one biological replicate (n = 5). Relative heat shock induction is statistically different in the strain expressing tRNA^Ser^_UGG,G26A_ compared to the control strain (Welch’s *t-*test; Bonferroni corrected *p*-value < 0.05).

We then performed an SGA analysis to identify genetic interactions with tRNA^Ser^_UGG,G26A_ using a collection of 1016 temperature sensitive alleles. The robotic screen identified 125 alleles with negative genetic interactions with tRNA^Ser^_UGG,G26A_. Genetic interactions were validated by remaking the double mutant strains, spotting normalized densities of the double mutants and their control strain on selective plates and measuring growth after two days (Supplemental File S2). After validation, 109 alleles representing 91 genes were classified as having a negative genetic interaction with tRNA^Ser^_UGG,G26A_ (Figure 2A).

**Figure 2.**
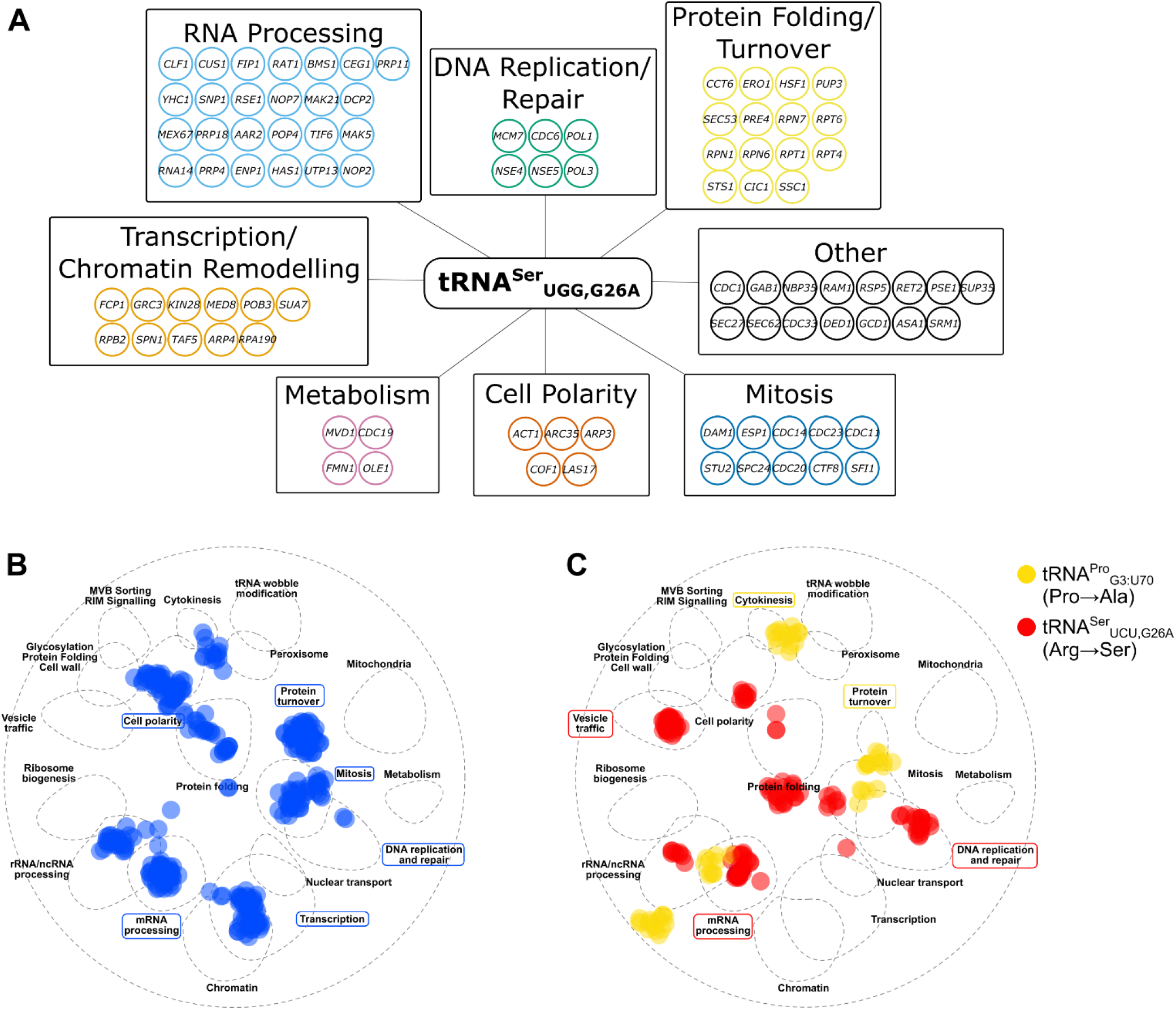
Negative genetic interaction network of the mistranslating tRNA^Ser^_UGG,G26A_. **A.** Genes validated as having a negative genetic interactions with tRNA^Ser^_UGG,G26A_ are arranged according to their predicted function based on gene descriptions in the yeast genome database (www.yeastgenome.org). **B.** SAFE analysis of genes that have a negative genetic interaction with tRNA^Ser^_UGG,G26A_ were mapped onto the yeast genetic interaction profile map (Costanzo *et al*. 2016) using TheCellMap (Usaj *et al*. 2017). Blue dots represent genes within the local neighborhood of genes validated to have negative genetic interactions with tRNA^Ser^_UGG,G26A_. Terms in blue boxes are network regions that are significantly enriched (Bonferroni corrected *p*-value < 0.05). **C.** SAFE analysis as performed in B with genetic interactions for tRNA^Pro^_G3:U70_ (yellow) and tRNA^Ser^_UCU,G26A_ (red) from Berg *et al*. (2021b). Terms in boxes represent network regions that are significantly enriched for the respectively mistranslating tRNA (Bonferroni corrected *p*-value < 0.05).

To further analyze the network of genes associated with the mistranslating tRNAs, we identified areas of the yeast genetic interaction map (Costanzo *et al*. 2016) that were enriched for negative genetic interactions with the mistranslating tRNA^Ser^_UGG,G26A_ using spatial analysis of functional enrichment (SAFE; Baryshnikova 2016; Figure 2B). Areas of the yeast genetic interaction network annotated with roles in protein turnover, cell polarity, mitosis, DNA replication and repair, transcription and mRNA processing were significantly enriched.

In a previous screen looking at negative synthetic genetic interactions with tRNA variants that mistranslate alanine at proline codons (tRNA^Pro^_G3:U70_; Hoffman *et al*. 2017a) and serine at arginine codons (tRNA^Ser^UCU,G26A; Berg *et al*. 2021b), we identified 10 and 47 negative genetic interactions, respectively. While these tRNA variants mistranslated at lower frequency (~ 3%) than tRNA^Ser^_UGG,G26A_ making specific comparison of genetic interactions difficult, it is possible to compare enriched pathways with a SAFE analysis (Figure 2C). tRNA^Ser^_UGG,G26A_ (Pro→Ser) and tRNA^Pro^_G3:U70_ (Pro→Ala) share negative genetic interactions enriched in the protein turnover area of the genetic network. tRNA^Ser^UCU,G26A (Arg→Ser) and tRNA^Ser^_UGG,G26A_ (Pro→Ser) interactions are enriched in DNA replication and repair and mRNA processing. Enrichment in cell polarity, mitosis and transcription were unique for genetic interactions with tRNA^Ser^_UGG,G26A_ (Pro→Ser) while enrichment in cytokinesis and vesicle trafficking were unique for tRNA^Pro^_G3:U70_ (Pro→Ala) and tRNA^Ser^UCU,G26A (Arg→Ser), respectively.

Only one strain had a positive genetic interaction with tRNA^Ser^_UGG,G26A_. The strain contains a temperature sensitive allele of *eco1*, an acetyltransferase required in sister chromatid cohesion. As we demonstrated previously (Zhu *et al*. 2020), the positive interaction results from tRNA^Ser^_UGG,G26A_ restoring serine at the S213P mutation of *eco1-1*.

### The frequency of mistranslation impacts the genetic interactions of tRNA^Ser^_UGG,G26A_

tRNA variants with the potential to mistranslate are found at numerous different loci in the human population (Berg *et al*. 2019a; Lant *et al*. 2019). Due to their ability to generate proteotoxic stress, we and others have suggested that these variants may be genetic modifiers of disease (Reverendo *et al*. 2014; Berg *et al*. 2017). In a previous analysis we demonstrated that, when comparing the same amino acid substitution, there is a near linear negative correlation between mistranslation frequency and cell growth in a wild-type *S. cerevisiae* background (Berg *et al*. 2019b). Our goal was to determine if the genetic background of strains having a synthetic interaction with tRNA^Ser^_UGG,G26A_ changes the impact of different mistranslation frequencies.

To regulate the frequency of mistranslation, we took advantage of our finding that placing a *GAL1* promoter (*GAL1pr*) sequence up or downstream of a tRNA gene represses tRNA expression when cells are grown in galactose (Berg *et al*. 2021a). Previously, using mass spectrometry, we determined the frequency of proline to serine mistranslation in strains containing centromeric plasmids expressing wild-type tRNA^Ser^, tRNA^Ser^_UGG,G26A_, tRNA^Ser^_UGG,G26A_ with 5’ *GAL1pr* and tRNA^Ser^_UGG, G26A_ with 3’ *GAL1pr* in galactose containing medium to be 0.3%, 5.6%, 3.5% and 0.9%, respectively (Figure 3A; Berg *et al*. 2021a). Three frequencies of mistranslation above background are thus achieved: the greatest mistranslation with no flanking *GAL1pr*, intermediate mistranslation with the *GAL1pr* upstream of the tRNA and the least mistranslation with the *GAL1pr* downstream of the tRNA.

**Figure 3.**
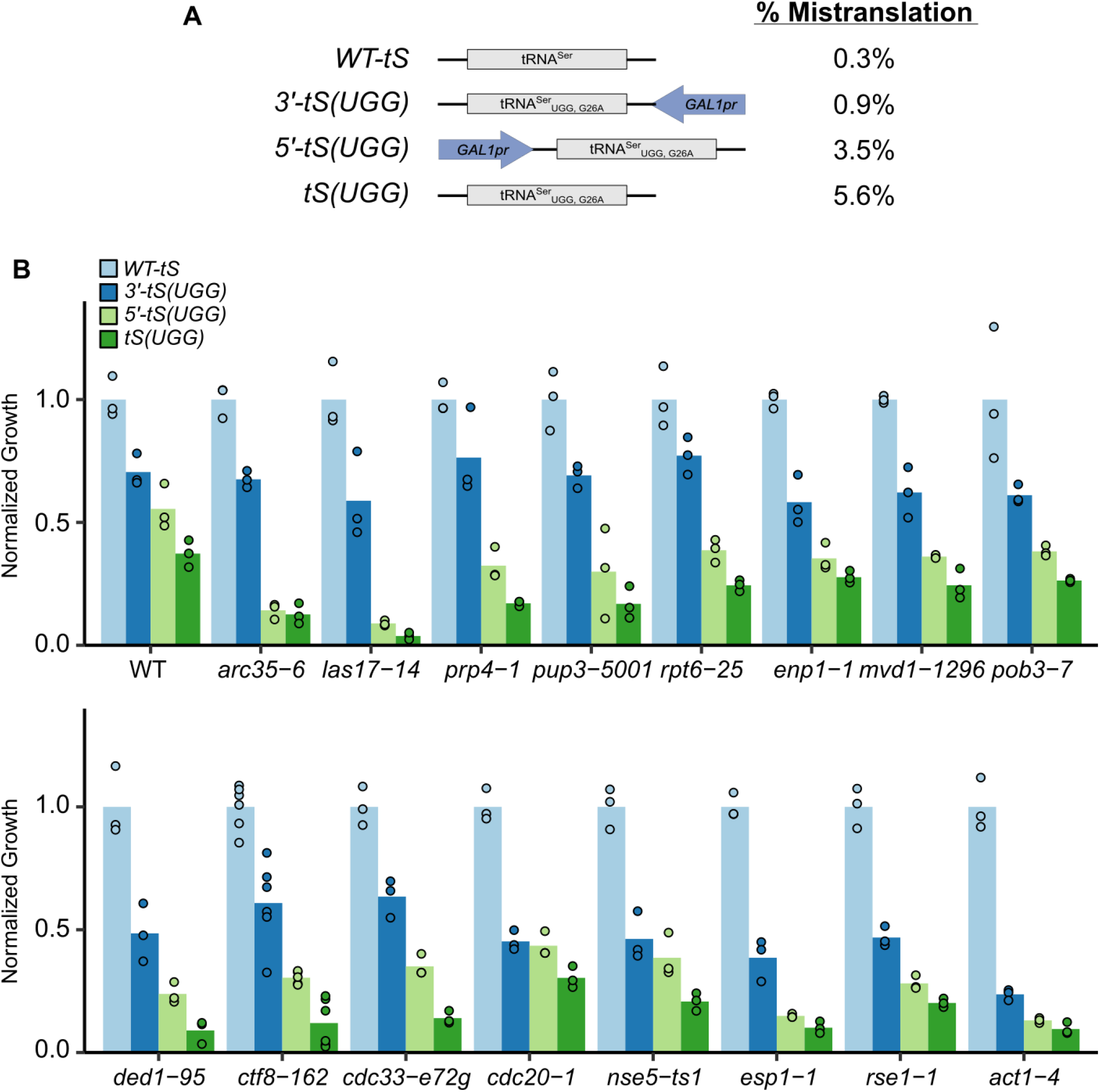
Effect of different mistranslation frequencies on growth differs depending on strain background. **A.** Schematic of the constructs containing wild-type tRNA^Ser^ [*WT-tS*], tRNA^Ser^_UGG,G26A_*-GAL1pr* [*3*’-*tS*(*UGG*)], *GAL1pr*-tRNA^Ser^_UGG,G26A_ [*5*’-*tS*(*UGG*)] and tRNA^Ser^_UGG,G26A_ [*tS*(*UGG*)] used to regulate proline to serine mistranslation frequency. Mistranslation frequencies were measured by mass spectrometry in Berg *et al*. (2021a). **B.** Wild-type BY4742 or the indicated strains from the temperature sensitive collection were transformed with the constructs described in A. Strains were grown to confluency in media lacking uracil and diluted 33-fold and spotted on media lacking uracil with galactose as the carbon source. The spot intensity of the strain containing the mistranslating tRNA was divided by the intensity of the strain containing the wild-type tRNA^Ser^ to determine normalized growth. Each point represents one biological replicate.

To determine the impact of mistranslation frequency, we selected 16 of the strains identified in the SGA analysis. These and the control strain were transformed with centromeric plasmids containing wild-type tRNA^Ser^, tRNA^Ser^_UGG, G26A_, 5’ *GAL1pr*-tRNA^Ser^_UGG, G26A_ and 3’ tRNA^Ser^_UGG, G26A_-*GAL1pr*. Triplicate cultures of independent transformants for each strain were grown to confluency, diluted 33-fold, spotted onto minimal medium with galactose as the carbon source and grown at 30°. Spot plates are shown in Figure S1. The density of the spotted cultures was measured and then expressed as a percentage of the spot density for the same strain background containing wild-type tRNA^Ser^. For example, tRNA^Ser^_UGG, G26A_ reduces growth of the wild-type BY4742 strain to 37 ± 5% of growth seen in BY4742 containing wild-type tRNA^Ser^. The 5’ *GAL1pr*-tRNA^Ser^_UGG, G26A_ and 3’ tRNA^Ser^_UGG, G26A_-*GAL1pr* reduce growth to 55 ± 9% and 71 ± 7% respectively.

As shown in Figure 3B and Figure S2, the wild type strain displays a near linear decrease with increasing mistranslation, consistent with our previous observations that increasing mistranslation frequency is negatively correlated with effects on growth for the same amino acid substitution (Berg *et al*. 2019b). All the synthetic strains showed a graded response where increased mistranslation results in more severe loss of growth, but interestingly, the pattern of decreased growth in response to changing the frequency of mistranslation differed amongst the strains. This difference suggests that the genetic background influences the impact of mistranslation.

To look at the patterns in more detail, we plotted the normalized growth of each temperature sensitive strain as a percentage of the growth of the wild-type strain (BY4742) for each of the three tRNA constructs that result in low, medium and high mistranslation frequency (Figure S3). We note that in these plots 100% indicates a lack of a negative synthetic effect, not the absence of an impact of mistranslation. Although the patterns appear to represent a continuum, we will focus the analysis on representative examples of four categories (Figure 4). In the first category are strains where the synthetic interaction with tRNA^Ser^_UGG,G26A_ increases proportionately with the frequency of mistranslation. In the representative examples *ctf8-62* and *cdc32-e72g* (Figure 4A), little synthetic interaction is seen at the lowest frequency of mistranslation. Many of the strains, including *arc35-6* and *las17-14* (Figure 4B) are in the second category. These show a modest synthetic effect at a low mistranslation frequency, but have a strong negative synthetic interaction at both moderate and high frequencies of mistranslation. The third category, represented best by *act1-4*, have a nearly equivalent synthetic effect at all three levels of mistranslation (Figure 4C). The last group includes *cdc20-1* and is related to group 3 but appears relatively more impacted by the lowest frequency of mistranslation (Figure 4D).

**Figure 4.**
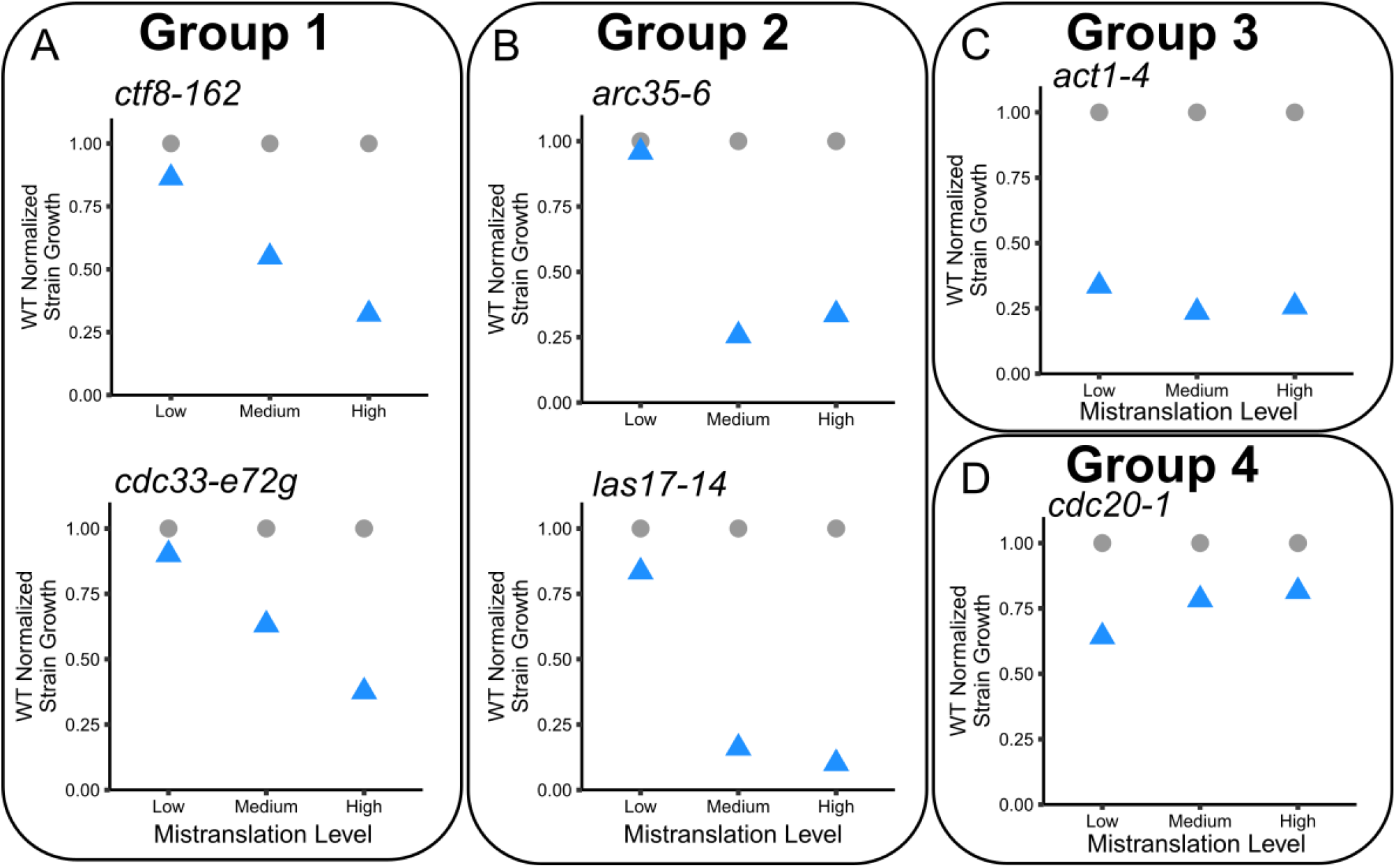
Genetic background alters the impact of different frequencies of proline to serine mistranslation. The average normalized growth calculated as in Figure 3 expressed as a percentage of the growth of the wild-type strain (BY4742) is shown in blue for the three different mistranslating constructs (Low: tRNA^Ser^_UGG,G26A_-*GAL1pr [3’-tS(UGG)]*, Medium: *GAL1pr*-tRNA^Ser^_UGG,G26A_ [*5*’-*tS*(*UGG*)] and High: tRNA^Ser^_UGG,G26A_ [*tS*(*UGG*)] for temperature sensitive strains expressing *ctf8-162* or *cdc33-e72g* **(A)**, *arc35-6* or *las17-14***(B),** *act1-4* **(C),** and *cdc20-1* **(D)**. The growth of the wild-type strain, 100%, is plotted as grey dots. Each point is the average of at least three biological replicates as in Figure 3.

Many factors determine the impact of a mistranslating tRNA. Factors intrinsic to the tRNA include the anticodon sequence and its resulting amino acid substitution (Berg *et al*. 2021b), the level of tRNA expression (Berg *et al*. 2021a) and the presence of secondary mutations that alter the stability of the tRNA (Berg *et al*. 2017, 2019b). Other factors are extrinsic to the tRNA. These include the number of competing tRNAs that buffer the mistranslating tRNA (Zimmerman *et al*. 2018), the environment in which cells expressing the mistranslating tRNA (Berg *et al*. 2021b) are found and the genetic background of the organism.

Genetic background contributes to the impact of a mistranslating tRNA in numerous ways. In yeast (this work and Hoffman *et al*. 2017a) and *Escherichia coli* (Ruan *et al*. 2008), loss of genes that regulate proteotoxic stress increase the severity of mistranslating tRNA variants, most likely because the tRNA variants increase the load of mis-made proteins. The impact of an extragenic mutation will depend on the extent to which it disrupts protein quality control or otherwise contributes to proteotoxic stress (Redler *et al*. 2016). Complexities arise since there are multiple quality control pathways that act independently but ultimately overlap to regulate proteostasis (reviewed in Chen *et al*. 2011). As such, impairing each pathway has the potential to show a different response to both changing levels of mistranslation and type of amino acid substitution. The nonlinear nature of the response likely arises because growth effects are not observed until a threshold of proteotoxicity is reached; different mutations will approach or exceed this threshold to different extents.

Other genetic mutations may exacerbate mistranslation if they occur in hypomorphic genes. Decreased protein level or function caused by mutation will be compounded by the reduced level of functional protein caused by mistranslation. The closer a genetic mutation brings the protein to the critical level of expression, the more impact the mistranslating tRNA will show when combined with that genetic mutant. Similar to the example above, until a critical threshold is exceeded, mistranslation may have little effect. At a more global level, genes that impact translation or mRNA processing can further limit expression of proteins already reduced by mistranslation.

Factors that alter the gene expression profile will also influence the impact of a mistranslating tRNA. Synonymous codon usage varies across the open reading frames in a genome (reviewed in Liu *et al*. 2021). Specific tRNA variants will mistranslate at a subset of these synonymous codons as determined by wobble rules and base modifications. tRNA variants only affect genes that are translated, and those that are more highly expressed will lead to greater proteotoxic stress when mistranslated. For yeast we have shown that one such factor is the environment in which the cells are grown (Berg *et al*. 2021b). As cell type determines gene expression profile in multicellular eukaryotes, different cell types are expected to be impacted differently by a tRNA variant. This argument is relevant to genetic background because mutations alter the internal and potentially external environment of the cell and often invoke a transcriptomic response (Hughes *et al*. 2000). The altered gene expression will in part determine the impact of a mistranslating tRNA.

The genetic background could also directly or indirectly alter expression of the mistranslating tRNA or that of competing endogenous tRNAs, altering the frequency of mistranslation. Genetic mutations that alter tRNA expression could occur in genes involved in regulating tRNA transcription (for example, by altering chromatin structure) or in genes with roles in tRNA processing, modification, nuclear import or stability. In our experiments, changes in expression of the mistranslating tRNA could occur if a secondary mutation occurs in a gene regulating transcription from the *GAL* promoter, though we note that none of the 16 temperature sensitive alleles we analyzed were in genes with annotated *GAL* regulatory roles.

## CONCLUSION

The impact of a mistranslating tRNA depends on a cell’s genetic background and the expression of the variant as it relates to the frequency of mistranslation. Approximately 20% of individuals contain a tRNA variant that potentially mistranslates (Berg *et al*. 2019a). The variants are found within different tRNA isodecoders at different loci, which is highly relevant since tRNA genes in the human genome are expressed at different levels (Torres *et al*. 2019). Our findings that genetic background influences the impact of different tRNA variants demonstrates that this must be taken into account when determining the contribution of tRNA variants to disease. Indeed, due to genetic or epigenetic differences, some individuals may be particularly sensitive to even low levels of mistranslation.

## ACKNOWLEDGMENTS

We would like to thank Onn Brandman for the GFP-Hsf1 reporter plasmid, Ricard Rodriguez-Mias for assisting with the mass spectrometry and maintaining the instruments and Ecaterina Cozma and Josh Isaacson for critically reading the manuscript.

